# Plasma metabolomics of an oral protein tolerance test reveals altered amino acid handling in sarcopenia

**DOI:** 10.64898/2026.06.29.735267

**Authors:** Tim Havers, Sebastian Martini, Maximillian Hillgärtner, Gautam Rana, Martin Schönfelder, Moritz Eggelbusch, Michael Witting, Dominik Lutter, Gülsah Erdogan, Karsten Köhler, Philipp Baumert, Stuart Phillips, Stephan Geisler, Michael Drey, Henning Wackerhage

## Abstract

**Background:** Sarcopenia is associated with anabolic resistance, a blunted muscle protein synthesis response to protein ingestion. Here, we hypothesized that anabolic resistance may be associated with a delayed postprandial decline in circulating plasma amino acids following protein ingestion. We therefore wanted to investigate whether an oral protein tolerance test (OPTT) combined with untargeted plasma metabolomics can detect age-related or sarcopenia-related differences in amino acid time courses consistent with altered postprandial amino acid handling, which could potentially reflect reduced anabolic sensitivity. Moreover, we investigated whether metabolites other than amino acids reacted to the OPTT.

**Methods:** Twelve young healthy adults (controls: 22-28 years) and 12 older adults with clinically diagnosed probable or confirmed sarcopenia (70-91 years) ingested 20 g of whey protein after an overnight fast. We collected venous blood at baseline, 1 h, and 2 h post-ingestion and analyzed the samples by untargeted LC-HRMS plasma metabolomics. Linear mixed-effects models were fitted for 2,968 metabolic features with Benjamini–Hochberg FDR correction. For each category (branched-chain amino acid, essential amino acid [EAA], total amino acid) we summed the within-subject log_₂_ fold changes (FC) of the constituent amino acids. This composite is reported as the summed log_₂_FC

**Results:** 201 metabolites were structurally annotated including 58 amino acid-related metabolites and 97 lipids. Fourteen of 17 proteinogenic amino acids increased significantly after protein ingestion (FDR<0.05). In young controls, essential amino acids rose more steeply at 1 h than in sarcopenic individuals (+10.06 ± 1.05 vs. +7.84 ± 1.58 summed log_₂_FC) and declined more between 1 and 2 h (−4.93 ± 1.29 vs. −0.20 ± 2.27 summed log_₂_FC). Leucine exemplified this pattern best, rising 1.74 log_₂_FC in controls and declining to 0.96 at 2 h, while remaining elevated at 1.61 log_₂_FC in the sarcopenic group at 2 h (p=0.009). Beyond amino acids, whey protein lowered circulating free fatty acids in both groups (FA 18:2, FA 18:1, FA 16:0; all FDR<0.05). Medium- and long-chain acylcarnitines (Car 8:0, Car14:2) declined postprandially in controls but remained elevated in sarcopenic individuals (p<0.05), suggesting altered postprandial lipid metabolism.

**Conclusion:** In this proof-of-concept study, an OPTT showed that plasma EAAs declined more slowly from their postprandial peak in older adults with sarcopenia than in young adults, consistent with altered postprandial amino acid handling that may reflect anabolic resistance. Whey protein ingestion additionally modulates lipid and acylcarnitine metabolism in an age-dependent manner, suggesting broader alterations in postprandial metabolic regulation in older adults with sarcopenia.

## Introduction

All ≈600 human skeletal muscles combined are the largest organ by mass in the human body, contributing on average 31% to body mass in women and 38% in men [1]. Sarcopenia is a progressive, generalised skeletal muscle disorder defined by low muscle strength together with low muscle quantity or quality, and, in severe cases, reduced physical performance [2]. It should be distinguished from the gradual, largely physiological loss of muscle mass that accompanies normal ageing (typically <1% per year in individuals over 60; [3]). Under the EWGSOP2 framework, sarcopenia denotes a clinical condition in which a pathological threshold is exceeded and that is associated with falls, frailty, loss of independence, and higher all-cause mortality [2]. Moreover, lower muscle mass is frequently accompanied by greater whole-body fat mass and poorer glucose homeostasis [4].

The pathogenesis of sarcopenia is multifactorial. One contributing mechanism is anabolic resistance, a blunted muscle protein synthesis response to anabolic stimuli.

Anabolic resistance refers to a reduced muscle protein synthesis response to an anabolic stimulus such as protein or essential amino acids [5, 6], resistance training [7], or pharmacologic agents [8]. Anabolic resistance is increasingly recognized as one of several mechanisms contributing to muscle loss in aging and disease [9]. Clinically, anabolic resistance may limit the effectiveness of interventions aimed at preserving or increasing muscle mass.

At present, anabolic sensitivity is commonly defined as the capacity of skeletal muscle to increase muscle protein synthesis in response to an anabolic stimulus, with anabolic resistance representing its impairment. It is typically assessed by applying an anabolic stimulus such as resistance exercise, protein ingestion, or anabolic hormone, and quantifying muscle protein synthesis through the incorporation of a stable isotope-labelled amino acid tracer into muscle protein. This approach requires tracer infusion and multiple muscle biopsies [10]. Although this is the gold standard, it is too invasive and too costly for clinical practice and thereby limited to research only. Currently, there is no proxy test for anabolic sensitivity, and so clinicians have no information about the anabolic sensitivity of their patients, preventing a personalization of the treatment of muscle wasting. Thus, the development of tests for anabolic sensitivity is a pressing clinical need.

We reasoned that an oral protein tolerance test (OPTT) might serve as a clinically feasible proxy test analogous to an oral glucose tolerance test (OGTT). In both cases, a substrate (i.e., protein versus glucose) is ingested which activates a polymer synthesis reaction: glucose drives glycogen synthesis, while protein, particularly leucine [11], stimulates muscle protein synthesis via activation of mTORC1. Muscle tissues take up circulating amino acids as a substrate for muscle protein synthesis. Therefore, the rate at which plasma amino acid levels decline after the postprandial peak may serve as an indirect proxy of whole-body postprandial elevations of amino acids, termed aminoacidaemia. We reasoned that if anabolic resistance impairs muscle protein synthesis in sarcopenic individuals, this postprandial decline should be attenuated compared with young adults. We acknowledge, however, that plasma amino acid concentration reflects a net balance between appearance, driven by gastric emptying, intestinal absorption, and splanchnic extraction, and disappearance across all tissues, including oxidation and non-muscle protein synthesis. Without stable isotope tracers, the specific contribution of muscle protein synthesis cannot be determined.

Another aspect of an oral protein tolerance test is that protein ingestion may also affect other metabolites via the activation of mTORC1. mTORC1, which is activated especially by leucine [11], stimulates not only protein synthesis but also autophagy, glycolysis, and nucleotide and lipid synthesis [12]. Therefore, increased mTORC1 activity after protein ingestion should also affect the concentrations of other metabolites, but this response is not well understood. However, it is important to characterize the metabolome response to protein ingestion, as an intake of protein greater than the recommended daily allowance is recommended e.g., for the older adults [13].

Based on this reasoning, we compared the plasma metabolome response to protein ingestion of young women and men to that of sarcopenic older adults. The aim of this experiment was to answer the following two research questions:

1. How do plasma amino acid levels respond to a protein bolus in older adults with sarcopenia versus young individuals, and is the response consistent with anabolic resistance in sarcopenic participants?
2. Do non-amino acid metabolites respond to the oral protein tolerance test, and are these responses influenced by age or sarcopenia?

## Methods

### Ethical approval

We obtained oral and written informed consent from all participants. The study adhered to the principles of the Declaration of Helsinki and was approved by the Ethics Committee of Ludwig Maximilian University of Munich, Germany (24-0250).

### Participants

We recruited 24 participants, including 12 young adults (18–30 years) and 12 older adults (70–90 years) with clinically diagnosed probable or confirmed sarcopenia (Table 1). Each group comprised six men and six women. Young individuals were eligible if they were 18–30 years old, were not obese, and, in the case of female participants, were not pregnant. Older adults were eligible if they were 70–90 years old, were not obese, and had a clinically diagnosed probable or confirmed sarcopenia during their visit to the Geriatric outpatient clinic. We diagnosed probable and confirmed sarcopenia according to the revised EWGSOP2 criteria (see supplementary material 1 for specifics) [2].

**Table 1:**
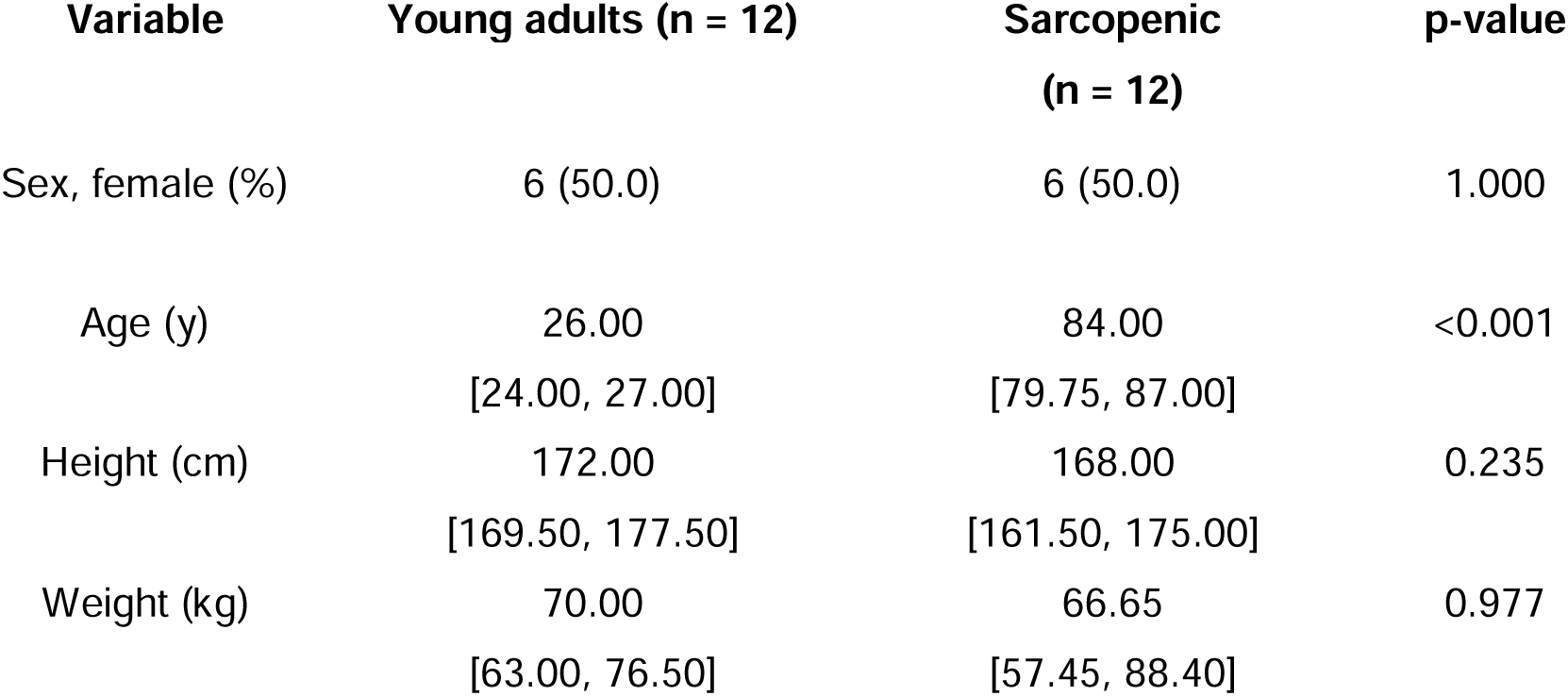
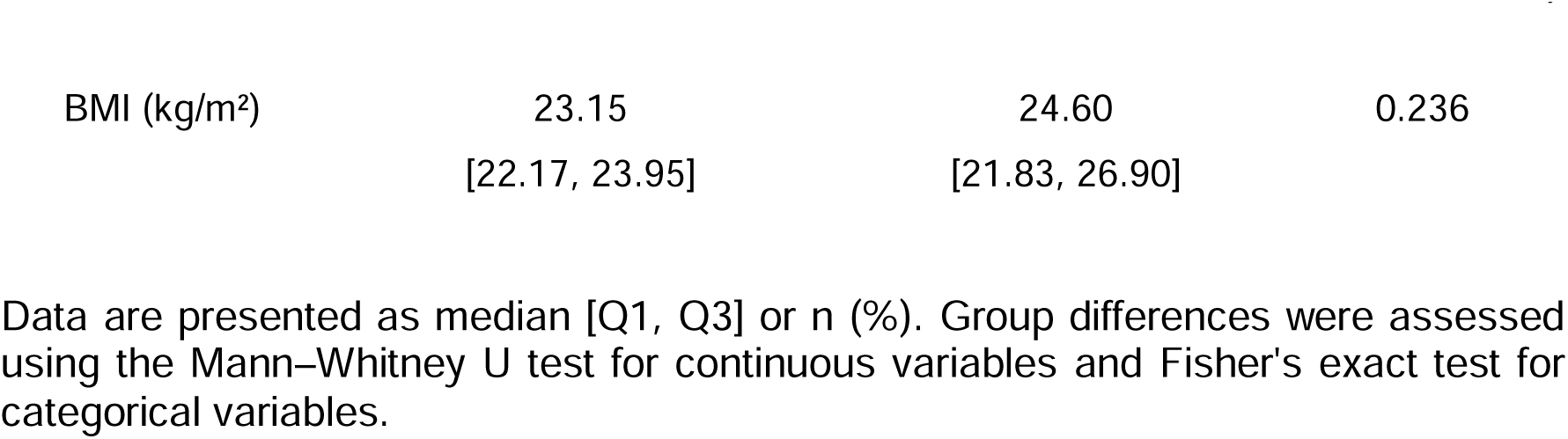
Demographic characteristics.

Exclusion criteria for both groups were residence in a nursing home, neuromuscular disorders (e.g., myasthenia gravis, muscular dystrophy, amyotrophic lateral sclerosis, poliomyelitis), moderate to severe dementia, chronic inflammatory conditions (e.g., Crohn’s disease, ulcerative colitis, rheumatoid arthritis), systemic corticosteroid therapy, a history of cancer within the past five years, and milk protein allergy or lactose intolerance.

### Study design

We conducted a monocentric, prospective, controlled, repeated-measures study (Figure 1a). All participants received a single standardised oral protein challenge: 23 g of whey protein (Fresubin® Protein Powder, Fresenius Kabi, neutral flavour) from a single batch dissolved in 200 mL water, providing 20.0 g protein and 82.8 kcal. The complete amino acid profile is shown in Table 2. We collected venous blood collected in the fasted state and at 1 h and 2 h post-ingestion. No randomization or blinding was applied. All were drawn within a 2 h time window.

**Figure 1.**
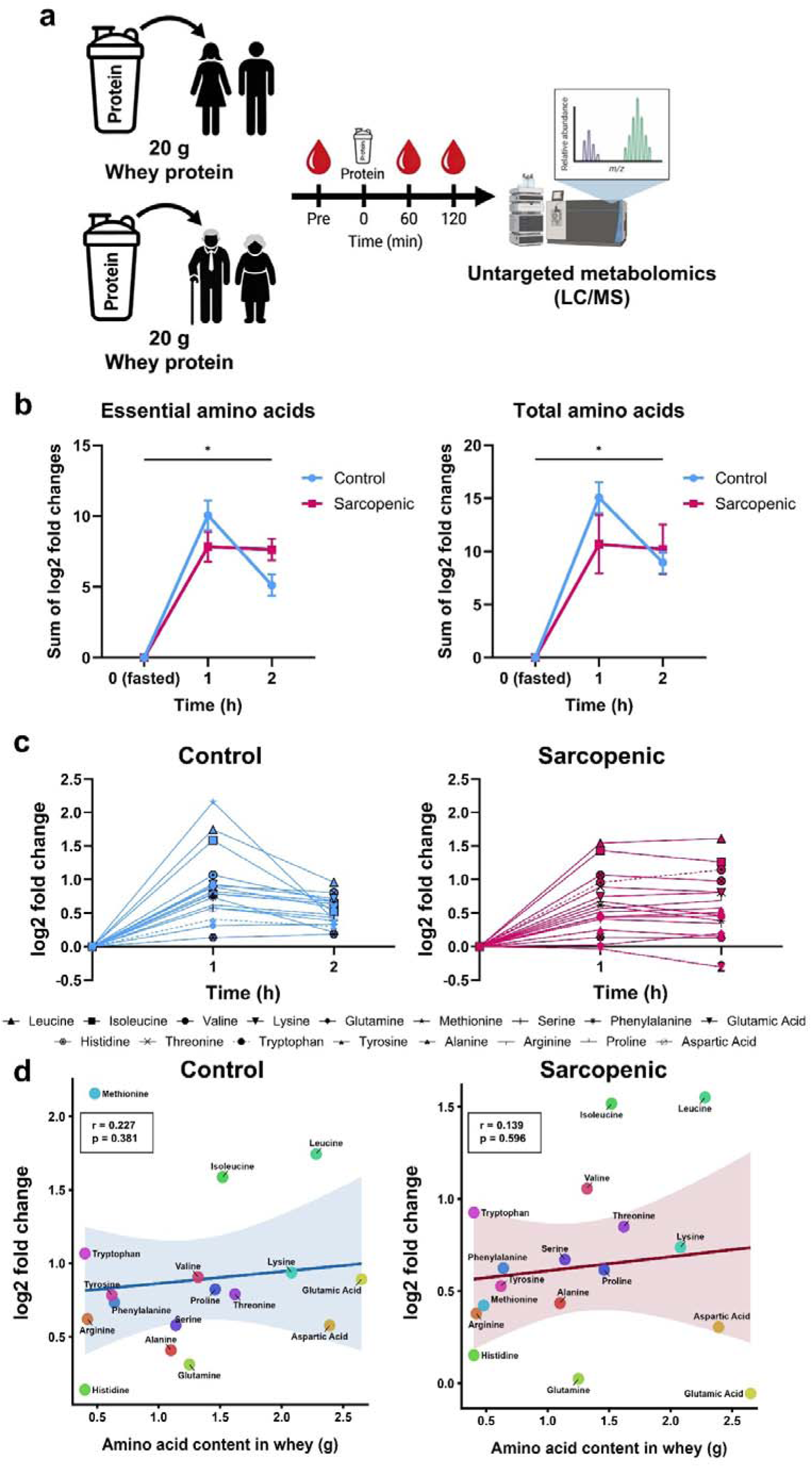
Study design, and amino acid response after protein ingestion. **a)** Study design. Sarcopenic and young healthy individuals received a whey protein bolus in the fasted state. At baseline, 1 h and 2 h after protein ingestion, venous blood samples were collected and processed to plasma and analyzed using untargeted metabolomics. **b)** Protein tolerance test expressed as the sum of log2 fold changes relative to baseline in essential amino acids and total amino acids in young control subjects compared with individuals with sarcopenia. Data are mean ± SEM *indicates a significant time × group interaction (linear mixed-effects model); main effects of time and group are reported in Supplementary Material 2 S-Table 5. **c)** Overview of the log2 fold change relative to baseline of proteinogenic amino acids in control subjects and individuals with sarcopenia, illustrating distinct postprandial trajectories. **d)** Spearman correlation analysis showing the relationship between amino acid content in whey protein and the log2-transformed fold change in plasma amino acid levels at 1 h post-ingestion. Solid lines represent linear regression fits, and shaded areas indicate 95% confidence intervals. No significant correlations were observed in either group.

**Table 2.** – Amino Acid Profile of the Fresubin® Protein Powder.

| Nutrient | Unit | per 100 g | per 23 g dose |
| --- | --- | --- | --- |
| Energy | kcal | 360 | 82.8 |
| Protein | g | 87.0 | 20.01 |
| Fat | g | 1.0 | 0.23 |
| Carbohydrates | g | ≤1.0 | ≤0.23 |
| ***Essential Amino Acids*** |  |  |  |
| Leucine | g | 9.92 | 2.28 |
| Lysine | g | 9.05 | 2.08 |
| Threonine | g | 7.05 | 1.62 |
| Isoleucine | g | 6.61 | 1.52 |
| Valine | g | 5.74 | 1.32 |
| Phenylalanine | g | 2.78 | 0.64 |
| Methionine | g | 2.09 | 0.48 |
| Tryptophan | g | 1.74 | 0.40 |
| ***Conditionally Essential Amino Acids*** |  |  |  |
| Glutamine | g | 5.43 | 1.25 |
| Tyrosine | g | 2.70 | 0.62 |
| Cysteine | g | 2.35 | 0.54 |
| Arginine | g | 1.83 | 0.42 |
| Histidine | g | 1.74 | 0.40 |
| ***Non-essential Amino Acids*** |  |  |  |
| Glutamic acid | g | 11.50 | 2.65 |
| Asparagine & Aspartic acid | g | 10.40 | 2.39 |
| Proline | g | 6.35 | 1.46 |
| Serine | g | 4.96 | 1.14 |
| Alanine | g | 4.79 | 1.10 |
| Glycine | g | 1.57 | 0.36 |

Values are taken from the manufacturer's product specification (Fresubin® Protein Powder, neutral flavour; Fresenius Kabi Deutschland GmbH, Bad Homburg, Germany).

### Anthropometric assessment

Stature and body weight were measured by using a standardized wall stadiometer (SECA 242, Seca, Hamburg, Germany) and a mechanical scale (platform scale SECA 635, Seca, Hamburg, Germany).

### Blood collection and processing

We collected venous blood from the antecubital vein into 4 mL EDTA tubes (fasted baseline, 60 min, 120 min), inverted 3–5 times, and centrifuged at 2,400 × g for 10 min at room temperature. We aspirated plasma without disturbing the buffy coat, transferred to cryovials, snap-frozen on dry ice, and stored at −80°C until analysis. We handled and stored all samples in pseudonymized form, and we did not use any stabilizing additives beyond standard EDTA.

### Metabolomics

All solvents and chemicals were of analytical grade or higher. We used only LC-MS grade solvents for all LC-MS step (see supplementary material 1 for specifics). Plasma samples were extracted as previously described [14]. For the HILIC separation, the LC-HRMS/MS measurements were performed using an Agilent 1290 Infinity II BioLC coupled to a Sciex ZenoTOF 7600 tandem mass spectrometer (MS/MS) equipped with an OptiFlow Turbo V electrospray ionization (ESI) source. Samples were analysed in both positive (Agilent Poroshell 120 HILIC-Z column) and negative (Waters Atlantis Premier BEH Z-HILIC column) ionization modes [14]. Processing of data was performed using mzmine 4.5 and included recalibration, peak picking and alignment, gap filling, isotope grouping as well as metabolite annotation [15]. Metabolite annotation was performed by matching against an in-house spectral database was well as multiple external database and using the in-silico tools Sirius and CSI:FingerID using hits with high COSMIC confidence scores [16–18]. Annotation confidence is reported in concordance with the Metabolomics Standard Initiative or the confidence levels described by Schymanski et al. [19, 20]. The confidence level of all metabolites reported in this study is 2 to 3 (see supplementary material 2 for specifics).

### Statistical analysis

All analyses were performed in R (v4.4.1). Metabolomics data were log_₁₀_-transformed, median-normalized, and batch-corrected using ComBat. Partial Least Squares Discriminant Analysis (PLS-DA) was performed with the mixOmics package.

Linear mixed-effects models (LMEs) were fitted for each of the 2,968 metabolic features using the lme4 and lmerTest packages, with the log_₁₀_-transformed abundance as the response variable. For each feature the model was:

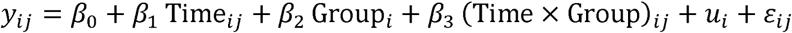

with *u_i_* ∼ *N*(0, 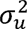) and *ε_ij_* ∼ *N*(0, *σ*^2^), where *y_ij_* is the log_₁₀_ abundance of participant *i* at time point *j*; Time (0, 1 h, 2 h), Group (young control, sarcopenic) and their interaction are fixed effects; *u_i_* is a participant-specific random intercept capturing between-subject variation in overall metabolite level, which thereby also accounts for the within-subject correlation induced by the repeated measurements; and *E_ij_* is the residual error. Models were estimated by maximum likelihood (REML = FALSE) using the bobyqa optimizer with a fallback to Nelder–Mead. Denominator degrees of freedom and p-values were approximated using the Satterthwaite method (the default in lmerTest). Type III F-tests were used to evaluate the main effects of Time and Group and their interaction. Multiple-testing correction was applied separately for each effect using the Benjamini–Hochberg false discovery rate (FDR) procedure across all 2,968 features.

To summarise the postprandial amino acid response, the abundance of each amino acid was first expressed as a within-subject log_₂_ fold change (FC) relative to that participant’s own fasting baseline.Because untargeted metabolomics yields relative signal intensities rather than absolute concentrations, these per-amino-acid log_₂_FCs were aggregated into a composite relative-response score for each category, branched-chain amino acids (leucine, isoleucine, valine), essential amino acids (branched-chain amino acids plus histidine, lysine, methionine, phenylalanine, threonine and tryptophan) and total amino acids (all proteinogenic amino acids present in the dataset), by summing the log_₂_FCs of the constituent amino acids at 1 h and at 2 h. This composite score reflects the cumulative relative change from baseline within a category and serves as an index of postprandial responsiveness.

Group differences in the composite amino acid response were assessed with separate LMEs for each category. For the composite score S of participant *i* at time point *j*,

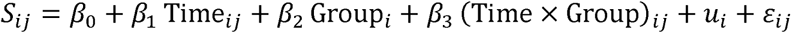

where Time (1 h, 2 h post-ingestion) was a within-subject fixed effect, Group (young controls, n = 12; sarcopenic older adults, n = 12) the between-subject fixed effect of primary interest, and *u_i_* a participant-specific random intercept. Models were fitted by maximum likelihood to enable likelihood-ratio-based inference. The Time × Group interaction was the term of interest, testing whether the change in the composite score from 1 h to 2 h differed between groups.

To assess whether the postprandial plasma response reflected the amino acid composition of the ingested bolus, the absolute amino acid content of the whey protein (g) was correlated with the individual log_₂_FC at 1 h using Spearman’s rank correlation, calculated separately for each group.

All visualisations were generated in R using ggplot2 (v3.4) and GraphPad Prism (v11.0.1). Statistical significance was assessed at two levels: a Benjamini–Hochberg-adjusted p-value < 0.05 to control the false discovery rate (FDR) across all 2,968 features, and a nominal p-value < 0.05 for exploratory, hypothesis-generating effects.

### Use of generative artificial intelligence

During the preparation of this manuscript, we used ChatGPT (OpenAI; GPT-5.5) solely to improve the language and readability of the text. No content, data, analyses, results, or interpretations were generated by this tool. The authors reviewed and edited all output and take full responsibility for the content of the publication.

## Results

We detected 2,968 metabolic features with the LC-MS method used, of which we structurally annotated 201 metabolites: 58 amino acids and related metabolites, 97 lipids, 9 carbohydrates, 9 nucleotides, 9 xenobiotics, 3 cofactors and vitamins, 2 energy metabolites, and 7 peptides. PLS-DA separated groups clearly across all metabolites (Component 1: 7.8%; Component 2: 6.0%) and across proteinogenic amino acids alone (Component 1: 53.7%; Component 2: 14.4%), confirming that sarcopenic individuals have a distinct basal plasma metabolome and respond differently to a protein ingestion than young individuals (Supplementary Material 2 Figure S1). We then used the annotated metabolomics data to answer our two research questions.

### How do plasma amino acid levels respond to a whey protein bolus in young versus sarcopenic women and men, and is the response consistent with anabolic resistance in sarcopenic participants?

Plasma amino acid responses to a 20 g whey protein bolus differed significantly between groups. In the composite linear mixed-effects model, the summed log_₂_ fold changes of branched-chain amino acids ((Supplementary Material 2: S-Table 5; S-Figure 3), essential amino acids, and total amino acids showed a significant time x group interaction (p < 0.05), indicating that the change in the postprandial composite score between 1 h and 2 h differed between young and sarcopenic participants. The sum of essential amino acids rose from baseline to 1 h after ingestion by 10.06 ± 1.05 log_₂_FC (mean ± SEM) in the young controls versus 7.84 ± 1.58 log_₂_FC in the sarcopenic older adults (Figure 1b). Consistent with our reasoning that a likely anabolic resistance would slow the postprandial decline in plasma amino acid abundance, the sum of essential amino acids declined by 4.93 ± 1.29 from 1 to 2 h but only by 0.20 ± 2.27 in the sarcopenic older adults. We observed a similar pattern for the sum of the log2-transformed levels of total amino acids (Figure 1b), suggesting a slower postprandial decline in plasma essential and total amino acid abundance from the 1-h peak following whey protein ingestion.

We next analyzed the proteinogenic amino acids individually (Figure 1e). The branched-chain amino acids isoleucine, leucine, valine, essential amino acids besides histidine, and non-essential amino acids apart from aspartic acid and glutamic acid, increased over time (Supplementary Material 2: S-Table 3; S-Figure 2). Leucine, methionine, and tryptophan abundance in young versus sarcopenic individuals differed from each other. In all cases, amino acid abundance rose at 1 h in both groups, while it declined in the young group and further rose or maintained their abundance at 2 h post whey protein ingestion in the sarcopenic group (Supplementary Material 2: S-Table 3; S-Figure 2).

To test whether the postprandial plasma response simply mirrored the amino acid composition of the bolus, we correlated the amino acid content of the whey protein (g per 20 g protein) with each amino acid’s log_₂_FC at 1 h, separately for each group (Spearman’s rank correlation). No significant relationship was found in either group (all p > 0.05; Figure 1d), indicating that the 1-h plasma response was not proportional to the ingested amount of each amino acid. This dissociation is expected, as plasma amino acid concentrations are governed not only by the ingested dose but also by absorption kinetics, first-pass splanchnic extraction, and endogenous amino acid turnover. The two extremes illustrate this: methionine produced one of the largest 1-h responses (log_₂_FC 2.16 in the young controls; Figure 1d) despite its low content in whey (0.48 g per 20 g protein), whereas glutamic acid, the most abundant amino acid in the bolus (2.65 g per 20 g protein), changed only marginally (log_₂_FC ≈ 0.06; Figure 1d), consistent with its extensive first-pass metabolism by the intestine, which limits its appearance in the systemic circulation.

### Do non-amino acid metabolites respond to the oral protein tolerance test, and are these responses influenced by age or sarcopenia?

Protein is digested into amino acids, so protein ingestion might be expected to cause merely a transient rise in plasma amino acids. However, especially leucine should activate mTORC1 [11] which in turn regulates autophagy, glycolysis, nucleotide and lipid synthesis [12]. We therefore also analyzed the plasma metabolome response beyond amino acids. Whey protein ingestion suppressed multiple free fatty acids, including linoleic acid (FA 18:2), oleic acid (FA 18:1), and palmitic acid (FA 16:0) (all FDR < 0.001). Significant postprandial changes were also observed for several lysophosphatidylcholines (e.g., LPC 9:1;O, LPC 22:6; FDR < 0.05) and acylcarnitines (e.g., Car 3:0, FDR = 0.004).

Hierarchical clustering revealed four distinct response patterns (Figure 2a):

1. Cluster 1: free fatty acids with stronger postprandial suppression in controls;
2. Cluster 2: acylcarnitines and carbohydrate metabolites with higher basal abundance in the sarcopenic group, comprising most of the species with a significant time × group interaction;
3. Cluster 3: a smaller mixed group with divergent trajectories between groups; and
4. Cluster 4: lysophosphatidylcholines, predominantly differing between groups.

**Figure 2.**
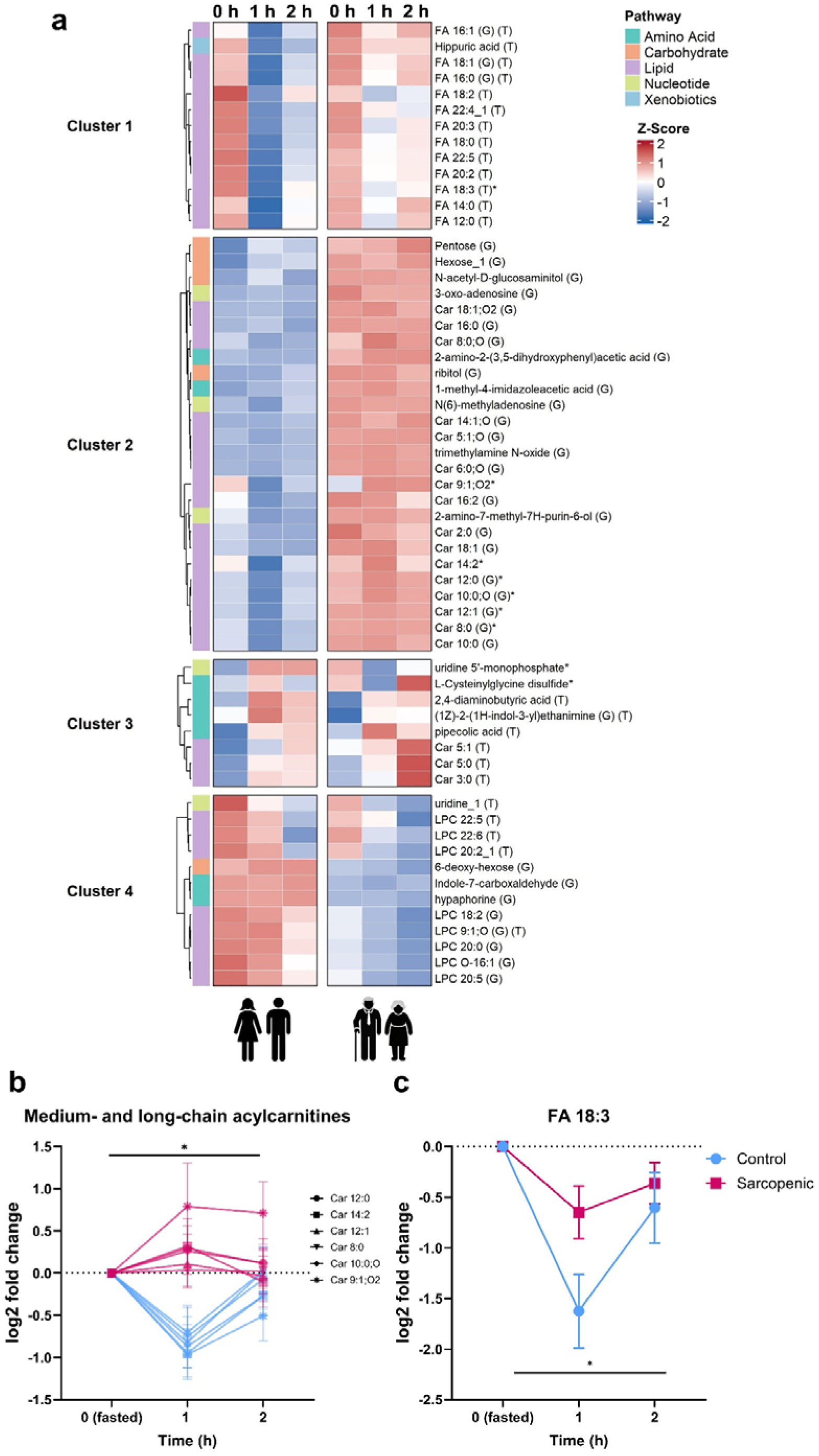
Global metabolic response to whey protein ingestion. **a)** Heatmap displaying significant metabolites (excluding proteinogenic amino acids) identified by FDR-corrected time or group effects, and/or nominally significant interaction effects (p < 0.05). Rows represent individual metabolites, while columns display the six experimental conditions (Control vs. Sarcopenic at 0h, 1 h, and 2 h). Data are shown as row Z-scores of log_₁₀_-transformed abundances. Hierarchical clustering (Ward’s linkage, Euclidean distance) identified three major response patterns: Cluster 1, free fatty acids showing a more pronounced postprandial reduction in the control group; Cluster 2, acylcarnitines and carbohydrate metabolites (e.g., pentoses, hexoses) with higher basal and postprandial abundances in the sarcopenic group, including most species with a significant time × group interaction; Cluster 3, a smaller mixed group of metabolites with divergent or opposing trajectories between groups; and Cluster 4, lysophosphatidylcholines, predominantly showing group differences. Row annotations denote metabolic super-pathways. Statistical markers: G = significant group differences (FDR corrected). T = significant time effects (FDR corrected). * = significant time x group interactions (nominal). **b-c)** Overview of the log2 fold changes relative to baseline of **(b)** medium- and long-chain acylcarnitines and **(c)** FA 18:3, illustrating distinct postprandial trajectories.

Several medium- and long-chain acylcarnitines showed divergent postprandial trajectories between groups (Figure 2b). In controls, Car 8:0, Car 10:0;O, and Car 14:2 declined at 1 h post-ingestion, whereas in the sarcopenic group these species remained elevated or increased (nominal interaction p < 0.05). Similarly, FA 18:3 dropped markedly at 1 h in controls but showed only a modest and transient decline in the sarcopenic group (Figure 2c). Uridine 5’-monophosphate and L-cysteinylglycine disulfide showed opposing trajectories between groups despite no overall time effect, with levels rising in one group while falling in the other.

## Discussion

Untargeted plasma metabolomics after a whey protein bolus revealed that total plasma amino acid levels declined more slowly after a 1 h peak in sarcopenic older adults than in young women and men. This altered postprandial amino acid handling is consistent with but does not prove anabolic resistance. In addition, protein ingestion not only altered amino acid levels but changed lipid- and acylcarnitine-related metabolite levels, with differences between sarcopenic older adults and young participants.

### Amino acid kinetics and implications for an oral protein tolerance test

Our first research question was whether postprandial amino acid kinetics are influenced by anabolic sensitivity. We found that total and essential amino acid levels declined less between 1 h and 2 h after whey protein ingestion in the sarcopenic older adults than in the young participants. This pattern is consistent with impaired peripheral amino acid utilization and anabolic resistance in older adults [5].

Among individual amino acids, leucine kinetics after whey ingestion differed most between the two groups. In young controls, leucine rose 1.74 log_₂_FC at 1 h and declined to 0.96 log_₂_FC at 2 h, consistent with greater net disappearance from plasma after the peak. In contrast, the sarcopenic group reached a similar peak of 1.55 log_₂_FC at 1 h but further increased to 1.61 log_₂_FC at 2 h, suggesting a slower postprandial decline in plasma leucine (nominal interaction effect: p < 0.05). This may be physiologically relevant because leucine is a key activator of mTORC1 signalling, which contributes to the regulation of muscle protein synthesis [11]. Leucine uptake into skeletal muscle depends partly on LAT1 (gene *SLC7A5*)-mediated transport [21], after which intracellular leucine activates mTORC1 via the Sestrin2–GATOR2 pathway [22]. The sustained elevation of circulating leucine in the older participants may therefore reflect a slower net disappearance from plasma, potentially consistent with reduced anabolic responsiveness, though contributions from altered splanchnic extraction, lower oxidation, or reduced whole-body protein turnover cannot be excluded without isotope tracer measurements.

Methionine and tryptophan also declined more slowly in the older participants than in the young. Methionine regulates mTORC1 signalling via SAMTOR [23]. Despite the low amount of methionine in whey, methionine abundance increased most in young participants, followed by a rapid decline towards baseline, whereas concentrations remained elevated in the sarcopenic participants. Altered tryptophan and kynurenine metabolism have also been linked to frailty, impaired muscle function, and skeletal muscle atrophy during ageing [24, 25]. Importantly, ageing alters splanchnic sequestration of dietary amino acids [26, 27] and so this could contribute to the findings.

### Response of metabolites other than amino acids

Our second research question was whether metabolites other than amino acids respond to protein ingestion. This is relevant because especially leucine activates mTORC1 [11], which is a universal regulator of metabolism that regulates not only protein synthesis but also autophagy, glycolysis, and lipid metabolism [12]. Consistent with our reasoning, whey protein ingestion altered several lipid- and acylcarnitine-related metabolites.

Free fatty acids declined after whey ingestion in both groups, consistent with insulin-mediated suppression of adipose tissue lipolysis following the well-characterized insulinotropic response to whey protein [28]. However, sarcopenic participants exhibited attenuated suppression. This pattern is unlikely to reflect a deficient insulinotropic response, as the postprandial insulin response following whey protein ingestion does not differ between older and younger adults [29]. The blunting therefore more plausibly reflects reduced sensitivity of adipose tissue to an adequate insulin signal, consistent with evidence that aging delays postprandial suppression of lipolysis even under comparable insulinemia [30]. Whether this attenuated FFA suppression is mechanistically linked to the anabolic resistance phenotype commonly observed in sarcopenia, or reflects a parallel feature of the same metabolic deterioration, cannot be determined from the present data and warrants targeted investigation.

Acylcarnitine profiles provided additional insight into metabolic flexibility. Young participants showed a transient decrease followed by recovery, consistent with physiological substrate switching between lipid and carbohydrate or amino acid oxidation [31]. In contrast, sarcopenic participants exhibited stable or elevated acylcarnitine levels, a pattern associated with incomplete β-oxidation and mitochondrial dysfunction [32].

Together, these findings suggest that the metabolic response to protein ingestion extends beyond amino acid handling and involves coordinated changes in lipid metabolism, insulin signalling, and mitochondrial substrate utilization.

### Implications of branched-chain amino acid kinetics for disease risk

The sum of branched-chain amino acids remained elevated for longer after the protein bolus in sarcopenic than in young participants (S-Figure 3). This matters because chronically elevated branched-chain amino acid concentrations associate with increased risk of type 2 diabetes [33], and cardiovascular disease [34]. However, Mendelian randomization studies consistently show that type 2 diabetes, insulin resistance, and obesity raise branched-chain amino acid levels, not the reverse, indicating that elevated branched-chain amino acids largely reflect underlying metabolic dysfunction rather than driving diabetes risk [35, 36]. The cardiovascular picture is more nuanced. Mendelian randomization studies may indicate a role for branched-chain amino acids in inducing hypertension and possibly ischemic heart disease, but not in myocardial infarction, stroke, or heart failure [37, 38]. Whether the transiently elevated postprandial branched-chain amino acids exposure we observed here carries long-term clinical relevance remains speculative and requires dedicated longitudinal study.

### Clinical implications

This study provides preliminary evidence that an oral protein tolerance test could potentially be developed into a clinically feasible test of anabolic sensitivity or at least of amino acid handling. Such a test could help clinicians individualize anabolic therapies, including testosterone or bimagrumab, according to the anabolic responsiveness of the treated patient. In contrast, the current gold standard requires stable isotope tracer infusion and repeated muscle biopsies, making it too invasive and too costly for routine clinical use.

However, several challenges must be addressed before such a test is used in the clinic. First, studies must determine whether amino acid kinetics during an oral protein tolerance test reflect the muscle protein synthesis response to the anabolic stimulus as a direct measure of anabolic sensitivity. Second, future work should identify the best suited stimulus (e.g., whey protein versus leucine) and whether dosing should be fixed or normalized to body weight. Third, additional studies are needed to define the most informative blood sampling time points, as the present study only measured metabolites at 1 h and 2 h after ingestion. Fourth, it remains unclear which analytes provide the best diagnostic information, ranging from leucine or essential amino acids alone or broader metabolomic profiling.

The principle of an oral protein tolerance test resembles that of an oral glucose tolerance test. In both cases, an ingested substrate is digested, absorbed, and taken up by tissues where it supports polymer synthesis, namely glycogen or protein synthesis. In addition, the substrate activates anabolic signaling pathways that promote this process: glucose primarily via insulin signaling and amino acids, especially leucine, via mTORC1 activation [11]. As with the oral glucose tolerance test, further studies are required to validate, refine, and simplify the oral protein tolerance test before clinical implementation.

### Strengths, limitations, and future directions

The main strength of this study is its breadth. By combining an oral protein tolerance test with untargeted metabolomics, we characterized the plasma metabolome response to a whey protein bolus across age and sarcopenia status.

We acknowledge several limitations. Most importantly, we did not directly measure anabolic sensitivity using the current gold-standard method, namely stable isotope-labelled amino acid tracer infusion combined with serial muscle biopsies to quantify muscle protein synthesis. However, this was an exploratory proof-of-concept study designed to test whether an oral protein tolerance test combined with metabolomics could detect differences in postprandial amino acid handling between young and sarcopenic individuals. Performing stable isotope tracer studies with repeated muscle biopsies in all 24 participants would have substantially increased cost, complexity, and participant burden, particularly in the old sarcopenic cohort. Moreover, anabolic resistance in older sarcopenic individuals has already been demonstrated convincingly in previous tracer-based studies [5]. Nevertheless, the present study cannot determine whether the observed amino acid kinetics directly reflect muscle protein synthesis, as postprandial amino acid concentrations are additionally influenced by gastric emptying, intestinal absorption, splanchnic extraction, amino acid oxidation, and renal handling [26, 27]. Concurrent stable isotope tracer measurements with arterio-venous balance or muscle biopsy are therefore still required to validate plasma amino acid kinetics as a proxy measure of anabolic sensitivity [39].

Second, the sample size was small (n = 24; n = 6 per sex per subgroup), limiting statistical power. Most group × time interactions were only nominally significant and require replication in larger cohorts. Third, age and sarcopenia status were completely co-linear in our design, as the young controls (median 26 years) and sarcopenic participants (median 84 years) differed by nearly six decades. Consequently, the observed differences may reflect biological ageing, comorbidities, inactivity, or polypharmacy in addition to sarcopenia.

Fourth, blood sampling was limited to 2 h and may therefore have missed the full return towards baseline in the older group. Finally, metabolite annotations below confidence level 1 [19] should be considered hypothesis-generating until confirmed by targeted analyses.

## Conclusion

A standardized oral protein tolerance test combined with untargeted metabolomics detected differences in postprandial amino acid handling consistent with, but not proof of, anabolic resistance in older sarcopenic adults. The oral protein tolerance test also revealed broader alterations in lipid and amino acid metabolism. Before this approach can be used clinically to assess anabolic competence, validation against concurrent stable isotope tracer-derived muscle protein synthesis measurements is required. Given the small sample size and the co-linearity of age and sarcopenia status, these findings should be considered hypothesis-generating and require replication in larger cohorts including age-matched non-sarcopenic controls.

## Acknowledgement

We thank all subjects that participated in this study.

## References

1. Janssen I, Heymsfield SB, Wang ZM, Ross R. Skeletal muscle mass and distribution in 468 men and women aged 18-88 yr. J Appl Physiol (1985). 2000;89:81–8. doi:10.1152/jappl.2000.89.1.81.

2. Cruz-Jentoft AJ, Bahat G, Bauer J, Boirie Y, Bruyère O, Cederholm T, et al. Sarcopenia: revised European consensus on definition and diagnosis. Age Ageing. 2019;48:16–31. doi:10.1093/ageing/afy169.

3. Mitchell WK, Williams J, Atherton P, Larvin M, Lund J, Narici M. Sarcopenia, dynapenia, and the impact of advancing age on human skeletal muscle size and strength; a quantitative review. Front Physiol. 2012;3:260. doi:10.3389/fphys.2012.00260.

4. Havers T, Held S, Schönfelder M, Geisler S, Wackerhage H. Effects of Skeletal Muscle Hypertrophy on Fat Mass and Glucose Homeostasis in Humans and Animals: A Narrative Review with Systematic Literature Search. Sports Med. 2025;55:1867–85. doi:10.1007/s40279-025-02263-w.

5. Cuthbertson D, Smith K, Babraj J, Leese G, Waddell T, Atherton P, et al. Anabolic signaling deficits underlie amino acid resistance of wasting, aging muscle. FASEB J. 2005;19:422–4. doi:10.1096/fj.04-2640fje.

6. Mitchell WK, Wilkinson DJ, Phillips BE, Lund JN, Smith K, Atherton PJ. Human Skeletal Muscle Protein Metabolism Responses to Amino Acid Nutrition. Adv Nutr. 2016;7:828S–38S. doi:10.3945/an.115.011650.

7. Kumar V, Selby A, Rankin D, Patel R, Atherton P, Hildebrandt W, et al. Age-related differences in the dose-response relationship of muscle protein synthesis to resistance exercise in young and old men. J Physiol. 2009;587:211–7. doi:10.1113/jphysiol.2008.164483.

8. Lach-Trifilieff E, Minetti GC, Sheppard K, Ibebunjo C, Feige JN, Hartmann S, et al. An antibody blocking activin type II receptors induces strong skeletal muscle hypertrophy and protects from atrophy. Mol Cell Biol. 2014;34:606–18. doi:10.1128/MCB.01307-13.

9. Burd NA, Gorissen SH, van Loon LJC. Anabolic resistance of muscle protein synthesis with aging. Exerc Sport Sci Rev. 2013;41:169–73. doi:10.1097/JES.0b013e318292f3d5.

10. Wilkinson DJ. Historical and contemporary stable isotope tracer approaches to studying mammalian protein metabolism. Mass Spectrom Rev. 2018;37:57–80. doi:10.1002/mas.21507.

11. Atherton PJ, Smith K, Etheridge T, Rankin D, Rennie MJ. Distinct anabolic signalling responses to amino acids in C2C12 skeletal muscle cells. Amino Acids. 2010;38:1533–9. doi:10.1007/s00726-009-0377-x.

12. Szwed A, Kim E, Jacinto E. Regulation and metabolic functions of mTORC1 and mTORC2. Physiol Rev. 2021;101:1371–426. doi:10.1152/physrev.00026.2020.

13. Bauer J, Biolo G, Cederholm T, Cesari M, Cruz-Jentoft AJ, Morley JE, et al. Evidence-based recommendations for optimal dietary protein intake in older people: a position paper from the PROT-AGE Study Group. J Am Med Dir Assoc. 2013;14:542–59. doi:10.1016/j.jamda.2013.05.021.

14. Artati A, Couacault P, Witting M. Nontargeted Metabolomics Using the Sciex ZenoTOF 7600. Methods Mol Biol. 2025;2925:1–23. doi:10.1007/978-1-0716-4534-5_1.

15. Heuckeroth S, Damiani T, Smirnov A, Mokshyna O, Brungs C, Korf A, et al. Reproducible mass spectrometry data processing and compound annotation in MZmine 3. Nat Protoc. 2024;19:2597–641. doi:10.1038/s41596-024-00996-y.

16. Dührkop K, Shen H, Meusel M, Rousu J, Böcker S. Searching molecular structure databases with tandem mass spectra using CSI:FingerID. Proc Natl Acad Sci U S A. 2015;112:12580–5. doi:10.1073/pnas.1509788112.

17. Dührkop K, Fleischauer M, Ludwig M, Aksenov AA, Melnik AV, Meusel M, et al. SIRIUS 4: a rapid tool for turning tandem mass spectra into metabolite structure information. Nat Methods. 2019;16:299–302. doi:10.1038/s41592-019-0344-8.

18. Hoffmann MA, Nothias L-F, Ludwig M, Fleischauer M, Gentry EC, Witting M, et al. High-confidence structural annotation of metabolites absent from spectral libraries. Nat Biotechnol. 2022;40:411–21. doi:10.1038/s41587-021-01045-9.

19. Schymanski EL, Jeon J, Gulde R, Fenner K, Ruff M, Singer HP, Hollender J. Identifying small molecules via high resolution mass spectrometry: communicating confidence. Environ Sci Technol. 2014;48:2097–8. doi:10.1021/es5002105.

20. Sumner LW, Amberg A, Barrett D, Beale MH, Beger R, Daykin CA, et al. Proposed minimum reporting standards for chemical analysis Chemical Analysis Working Group (CAWG) Metabolomics Standards Initiative (MSI). Metabolomics. 2007;3:211–21. doi:10.1007/s11306-007-0082-2.

21. Hodson N, Brown T, Joanisse S, Aguirre N, West DWD, Moore DR, et al. Characterisation of L-Type Amino Acid Transporter 1 (LAT1) Expression in Human Skeletal Muscle by Immunofluorescent Microscopy. Nutrients 2017. doi:10.3390/nu10010023.

22. Saxton RA, Sabatini DM. mTOR Signaling in Growth, Metabolism, and Disease. Cell. 2017;168:960–76. doi:10.1016/j.cell.2017.02.004.

23. Gu X, Orozco JM, Saxton RA, Condon KJ, Liu GY, Krawczyk PA, et al. SAMTOR is an S-adenosylmethionine sensor for the mTORC1 pathway. Science. 2017;358:813–8. doi:10.1126/science.aao3265.

24. Al Saedi A, Chow S, Vogrin S, Guillemin GJ, Duque G. Association Between Tryptophan Metabolites, Physical Performance, and Frailty in Older Persons. Int J Tryptophan Res. 2022;15:11786469211069951. doi:10.1177/11786469211069951.

25. Kaiser H, Yu K, Pandya C, Mendhe B, Isales CM, McGee-Lawrence ME, et al. Kynurenine, a Tryptophan Metabolite That Increases with Age, Induces Muscle Atrophy and Lipid Peroxidation. Oxid Med Cell Longev. 2019;2019:9894238. doi:10.1155/2019/9894238.

26. Trommelen J, Holwerda AM, Pinckaers PJM, van Loon LJC. Comprehensive assessment of post-prandial protein handling by the application of intrinsically labelled protein in vivo in human subjects. Proc Nutr Soc. 2021;80:221–9. doi:10.1017/S0029665120008034.

27. Volpi E, Mittendorfer B, Wolf SE, Wolfe RR. Oral amino acids stimulate muscle protein anabolism in the elderly despite higher first-pass splanchnic extraction. Am J Physiol. 1999;277:E513–20. doi:10.1152/ajpendo.1999.277.3.E513.

28. Yanagisawa Y. How dietary amino acids and high protein diets influence insulin secretion. Physiol Rep. 2023;11:e15577. doi:10.14814/phy2.15577.

29. Giezenaar C, Hutchison AT, Luscombe-Marsh ND, Chapman I, Horowitz M, Soenen S. Effect of Age on Blood Glucose and Plasma Insulin, Glucagon, Ghrelin, CCK, GIP, and GLP-1 Responses to Whey Protein Ingestion. Nutrients 2017. doi:10.3390/nu10010002.

30. Osmond AD, Leija RG, Arevalo JA, Curl CC, Duong JJ, Huie MJ, et al. Aging delays the suppression of lipolysis and fatty acid oxidation in the postprandial period. J Appl Physiol (1985). 2024;137:1200–19. doi:10.1152/japplphysiol.00437.2024.

31. Kelley DE, Mandarino LJ. Fuel selection in human skeletal muscle in insulin resistance: a reexamination. Diabetes. 2000;49:677–83. doi:10.2337/diabetes.49.5.677.

32. Muoio DM. Metabolic inflexibility: when mitochondrial indecision leads to metabolic gridlock. Cell. 2014;159:1253–62. doi:10.1016/j.cell.2014.11.034.

33. Wang TJ, Larson MG, Vasan RS, Cheng S, Rhee EP, McCabe E, et al. Metabolite profiles and the risk of developing diabetes. Nat Med. 2011;17:448–53. doi:10.1038/nm.2307.

34. Murashige D, Jung JW, Neinast MD, Levin MG, Chu Q, Lambert JP, et al. Extra-cardiac BCAA catabolism lowers blood pressure and protects from heart failure. Cell Metab. 2022;34:1749–1764.e7. doi:10.1016/j.cmet.2022.09.008.

35. Mosley JD, Shi M, Agamasu D, Vaitinadin NS, Murthy VL, Shah RV, et al. Branched-chain amino acids and type 2 diabetes: a bidirectional Mendelian randomization analysis. Obesity (Silver Spring). 2024;32:423–35. doi:10.1002/oby.23951.

36. Doestzada M, Zhernakova DV, van den C L Munckhof I, Wang D, Kurilshikov A, Chen L, et al. Systematic analysis of relationships between plasma branched-chain amino acid concentrations and cardiometabolic parameters: an association and Mendelian randomization study. BMC Med. 2022;20:485. doi:10.1186/s12916-022-02688-4.

37. Zuo Z, Tong Y, Li M, Wang Z, Wang X, Guo X, et al. Effect of genetically determined BCAA levels on cardiovascular diseases and their risk factors: A Mendelian randomization study. Nutr Metab Cardiovasc Dis. 2023;33:2406–12. doi:10.1016/j.numecd.2023.08.003.

38. Zhao JV, Fan B, Burgess S. Using genetics to examine the overall and sex-specific associations of branch-chain amino acids and the valine metabolite, 3-hydroxyisobutyrate, with ischemic heart disease and diabetes: A two-sample Mendelian randomization study. Atherosclerosis. 2023;381:117246. doi:10.1016/j.atherosclerosis.2023.117246.

39. Mazzulla M, Hodson N, Scaife PJ, Smith K, Atherton PJ, Burd NA, et al. Oral amino acid tracer delivery detects feeding and exercise changes in myofibrillar protein synthesis rates in male adults. Physiol Rep. 2026;14:e70776. doi:10.14814/phy2.70776.

