## Supplementary Material 1 for "Plasma metabolomics of an oral protein tolerance test reveals altered amino acid handling in sarcopenia"

*shared first and presenting authors

**Supplementary Methods S1. Diagnosis of sarcopenia**

We diagnosed probable and confirmed sarcopenia according to the revised EWGSOP2 criteria (Cruz-Jentoft et al., 2019). Low muscle strength was assessed by handgrip dynamometry (JAMAR, Los Angeles, CA) and the five-times chair-stand test, and low muscle quantity by appendicular skeletal muscle mass index derived from dual-energy X-ray absorptiometry (Lunar Prodigy, GE Healthcare Technologies, USA). Participants with low muscle strength alone were classified as having probable sarcopenia, and those with low strength plus low appendicular skeletal muscle mass index as having confirmed sarcopenia. Of the 12 older participants, 7 met the criteria for probable and 5 for confirmed sarcopenia (Table 2).

**Supplementary Methods 1 - Table 1. Functional and body-composition characteristics of the older participants by EWGSOP2 sarcopenia status.**

| **Variable** | **Probable sarcopenia (n = 7)** | **Confirmed sarcopenia (n = 5)** |
| --- | --- | --- |
| Age, years | 87.0 [84.0–88.5] | 81.0 [76.0–83.0] |
| Sex, male/female | 3/4 | 3/2 |
| BMI, kg/m² | 26.0 [23.5–29.0] | 24.0 [20.0–25.0] |
| Chair-rise test completed, n/N (%) | 3/7 (42.9) | 2/5 (40.0) |
| Chair-rise time, s* | 20.8 [17.4–20.9] | 15.6 [15.2–16.0] |
| Grip strength, kg | 25.0 [24.4–28.2] | 20.8 [20.4–24.3] |
| ASMI, kg/m² | 7.04 [6.00–7.30] | 5.80 [5.40–6.14] |

Values are median [Q1–Q3] unless otherwise indicated. Probable sarcopenia denotes low muscle strength (EWGSOP2 Criterion 1); confirmed sarcopenia denotes low strength plus low appendicular skeletal muscle mass index (ASMI) by DXA (Criterion 2) (Cruz-Jentoft et al., 2019). *Chair-rise time refers only to participants able to complete the five-times chair-stand test; inability to complete the test owing to insufficient strength was classified as poor physical performance and reported as test not completed. ASMI, appendicular skeletal muscle mass index; DXA, dual-energy X-ray absorptiometry.

**Supplementary Methods S2. Metabolite extraction and LC-HRMS/MS parameters**

Plasma samples were extracted as previously described (Artati et al., 2025). Briefly, frozen plasma was thawed on ice and a pooled QC sample was prepared by combining aliquots from each sample; a commercial plasma sample served as long-term reference. For extraction, 100 µL of plasma or QC sample was mixed with 500 µL extraction solvent (MeOH with internal standards) and shaken at 1200 rpm for 2 min. After centrifugation at 2650 rpm, 50 µL aliquots of the supernatant were transferred to each of four 96-well plates and evaporated to dryness under nitrogen using a TurboVap 96 (Biotage). Plates were sealed and extracts stored dry at −80 °C until analysis.

Prior to measurement, two aliquots of dried extract were reconstituted with LC-compatible solvents containing chemical standards at fixed concentrations. The first aliquot was analysed in positive ionization mode on an Agilent Infinity Poroshell 120 HILIC-Z column (100 mm × 2.1 mm ID, 2.7 µm, PEEK-lined) with a gradient from eluent A (100 % H₂O + 10 mM ammonium formate / 0.1 % formic acid) to eluent B (10 % H₂O / 90 % ACN + 10 mM ammonium formate / 0.1 % formic acid). The second aliquot was analysed in negative ionization mode on a Waters Atlantis Premier BEH Z-HILIC column (100 mm × 2.1 mm ID, 1.7 µm) with a gradient from eluent A (100 % water + 10 mM ammonium acetate / ammonium hydroxide, pH 9) to eluent B (10 % water / 90 % acetonitrile + 10 mM ammonium acetate / ammonium hydroxide, pH 9). For both methods, flow rate was 0.5 mL/min, column temperature 40 °C, and injection volume 5 µL. Detailed LC and MS parameters are given in Artati et al. (2025).

Initial data QC comprised checking internal standards for retention time, m/z and intensity; all internal standards were detected with a mass error of ≤5 ppm.

**References:**

Artati, A., Couacault, P., & Witting, M. (2025). Nontargeted Metabolomics Using the Sciex ZenoTOF 7600. *Methods in Molecular Biology (Clifton, N.J.)*, *2925*, 1–23. <https://doi.org/10.1007/978-1-0716-4534-5_1>

Cruz-Jentoft, A. J., Bahat, G., Bauer, J., Boirie, Y., Bruyère, O., Cederholm, T., Cooper, C., Landi, F., Rolland, Y., Sayer, A. A., Schneider, S. M., Sieber, C. C., Topinkova, E., Vandewoude, M., Visser, M., & Zamboni, M. (2019). Sarcopenia: Revised European consensus on definition and diagnosis. *Age and Ageing*, *48*(1), 16–31. https://doi.org/10.1093/ageing/afy169
